# Viral RNA-dependent RNA polymerase inhibitor 7-deaza-2′-*C*-methyladenosine prevents death in a mouse model of West Nile virus infection

**DOI:** 10.1101/432955

**Authors:** Luděk Eyer, Martina Fojtíková, Radim Nencka, Ivo Rudolf, Zdeněk Hubálek, Daniel Ruzek

## Abstract

West Nile virus (WNV) is a medically important emerging arbovirus causing serious neuroinfections in humans against which no approved antiviral therapy is currently available. In this study, we demonstrate that 2′-*C*- methyl- or 4′-azido-modified nucleosides are highly effective inhibitors of WNV replication, showing nanomolar or low micromolar anti-WNV activity and negligible cytotoxicity in cell culture. One representative of *C*2′-methylated nucleosides, 7-deaza-2′-*C*- methyladenosine, significantly protected WNV-infected mice from disease progression and mortality. Twice daily treatment at 25 mg/kg starting at the time of infection resulted in 100% survival of the mice. This compound was highly effective, even if the treatment was initiated 3 days post-infection, at the time of a peak of viremia, which resulted in a 90% survival rate. However, the antiviral effect of 7-deaza-2′-*C*- methyladenosine was absent or negligible when the treatment was started 8 days post-infection (i.e., at the time of extensive brain infection). The 4′-azido moiety appears to be another important determinant for highly efficient inhibition of WNV replication in vitro. However, the strong anti-WNV effect of 4′-azidocytidine and 4′-azido-aracytidine was cell type-dependent and observed predominantly in PS cells. The effect was much less pronounced in Vero cells. Our results indicate that 2′-*C*- methylated or 4′-azidated nucleosides merit further investigation as potential therapeutic agents for treating WNV infections, as well as infections caused by other medically important flaviviruses.

## Introduction

West Nile virus (WNV) is an emerging representative of the genus *Flavivirus* belonging to the family *Flaviviridae*. This family also includes other medically important human pathogens, such as Dengue virus (DENV), Japanese encephalitis virus (JEV), yellow fever virus (YFV), Zika virus (ZIKV), and tick-borne encephalitis virus (TBEV) (1). WNV has been serologically classified into the JEV antigenic complex and divided into eight genotypic lineages; from a medical point of view, the strains pathogenic for humans are designated as lineages 1 and 2 (2). WNV virions (50 nm diameter) are enveloped with a host cell-derived lipid bilayer containing single-stranded, plus-sense genomic RNA 11 kb in length. The WNV genome encodes a single polyprotein processed into three structural (capsid, premembrane or membrane, and envelope) and seven nonstructural proteins (NS1, NS2A, NS2B, NS3, NS4A, NS4B, and NS5) (3–6). NS5 shows RNA-dependent RNA polymerase (RdRp) activity and has been reported to be an important target for antiviral development (4).

WNV circulates in nature within an enzootic transmission cycle between birds as reservoir hosts and bird-feeding mosquitoes, primarily involving *Culex* spp. as principal vectors of WNV (7). WNV was originally isolated in Africa in 1937 (8), and later caused outbreaks in Europe, the Middle East, and parts of Asia and Australia (9). Following its introduction into the United States in 1999, WNV rapidly disseminated across North America and has further spread to Mexico, South America, and the Caribbean (10,11). In humans, WNV infection often remains subclinical (12,13), but 20 – 40% of those infected may develop WNV disease, which manifests as a febrile illness that can progress to lethal encephalitis. Symptoms include cognitive dysfunction and acute flaccid paralysis (14–16). Overall, 2,002 cases of WNV disease in humans were reported in the United States in 2017, 67% of which were classified as neuroinvasive disease (e.g., meningitis or encephalitis) and 33% as non- neuroinvasive infection (https://www.cdc.gov/westnile). As no vaccines or specific therapies for WNV are currently approved for humans, there is an urgent need for an effective approach to treatment based on specific inhibitors of WNV replication (17).

Nucleoside analogs represent an important group of small molecule-based inhibitors that have figured prominently in the search for effective antiviral agents (18). After they enter the cell, nucleoside analogs are phosphorylated by cellular kinases and incorporated into viral nascent RNA chains (19,20). As the 3′-hydroxyl of nucleoside inhibitors is conformationally constrained or sterically/electronically hindered (“non-obligate chain terminators“), or completely missing (“obligate chain terminators“), such structures exert the decreased potency to form a phosphodiester linkage with the incoming nucleoside triphosphate during viral RNA replication, resulting in the premature termination of viral nucleic acid synthesis (21). Currently, there are more than 25 approved therapeutic nucleosides used for the treatment of viral infections of high medical importance, such as HIV/AIDS, hepatitis B, hepatitis C, and herpes infections (22–26). Therefore, nucleoside analogs represent promising tools that can be repurposed against mosquito-transmitted flaviviruses, including WNV.

The aim of this study was to assess anti-WNV activity and cytotoxicity in a series of methyl- or azido-substituted nucleosides. We have demonstrated that the 2′-*C*- methyl or 4′-azido substituents are crucial structural elements for highly efficient WNV inhibition and low cytotoxicity in cell culture. One representative of 2′methylated nucleosides, 7-deaza-2′-*C*- methyladenosine (7-deaza-2′-CMA), significantly protected WNV-infected mice from disease progression and mortality, even if the treatment was started 3 days post-infection (p.i.). Thus, 2′-*C*- methylated or 4′-azidated nucleosides merit further investigation as therapeutic agents for treating WNV infections and, hypothetically, infections caused by other medically important flaviviruses.

## Results

### In vitro antiviral effect and cytotoxicity of the tested compounds

Nucleosides with a methyl substituent at the *C*2′position exhibited nanomolar or low-micromolar dose-dependent anti-WNV activity when tested on either PS or Vero cell cultures (Table 1, Fig. 1A,B, and Fig. 2A,B). 7-Deaza-2′-CMA, also denoted as MK-0608, exerted the strongest antiviral effect of all 2′-*C*- methylated nucleosides tested, characterized by EC_50_ values of 0.33 ± 0.08 µM and 0.15 ± 0.05 µM for Eg-101 and 13–104, respectively. EC_50_ values for 2′-*C*- methyladenosine, 2′-*C*- methylguanosine, and 2′-*C*- methylcytidine were slightly higher compared to those for 7-deaza-2′-CMA, ranging from 0.78 ± 0.16 µM to 0.96 ± 0.06 µM for Eg-101, and 0.66 ± 0.15 µM to 1.67 ± 0.50 µM for 13–104. Similar to BCX-4430, which is used as a positive control for in vitro antiviral screens, these compounds (at a concentration of 50 µM) decreased the viral titer 10^5^ to 10^6^-fold compared to mock-treated cells, and the in vitro antiviral effect was stable up to 5 days p.i. (Fig. 2C). 2′-*C*-Methyluridine was almost inactive against both WNV strains (EC_50_ > 50 µM) when tested on porcine kidney stable (PS) cell monolayers, and exhibited only weak inhibitory activity in Vero cells (Fig. 1A,B). 2′-*C*-Methyl-modified nucleosides exerted low cytotoxicity in both PS and Vero cells (CC_50_ > 50 µM) (Table 1, Fig 1C), except 2′-*C*- methylcytidin had a CC_50_ value of ∼50 µM in PS 3 days post-treatment, as we reported previously (27).

**Figure 1.**
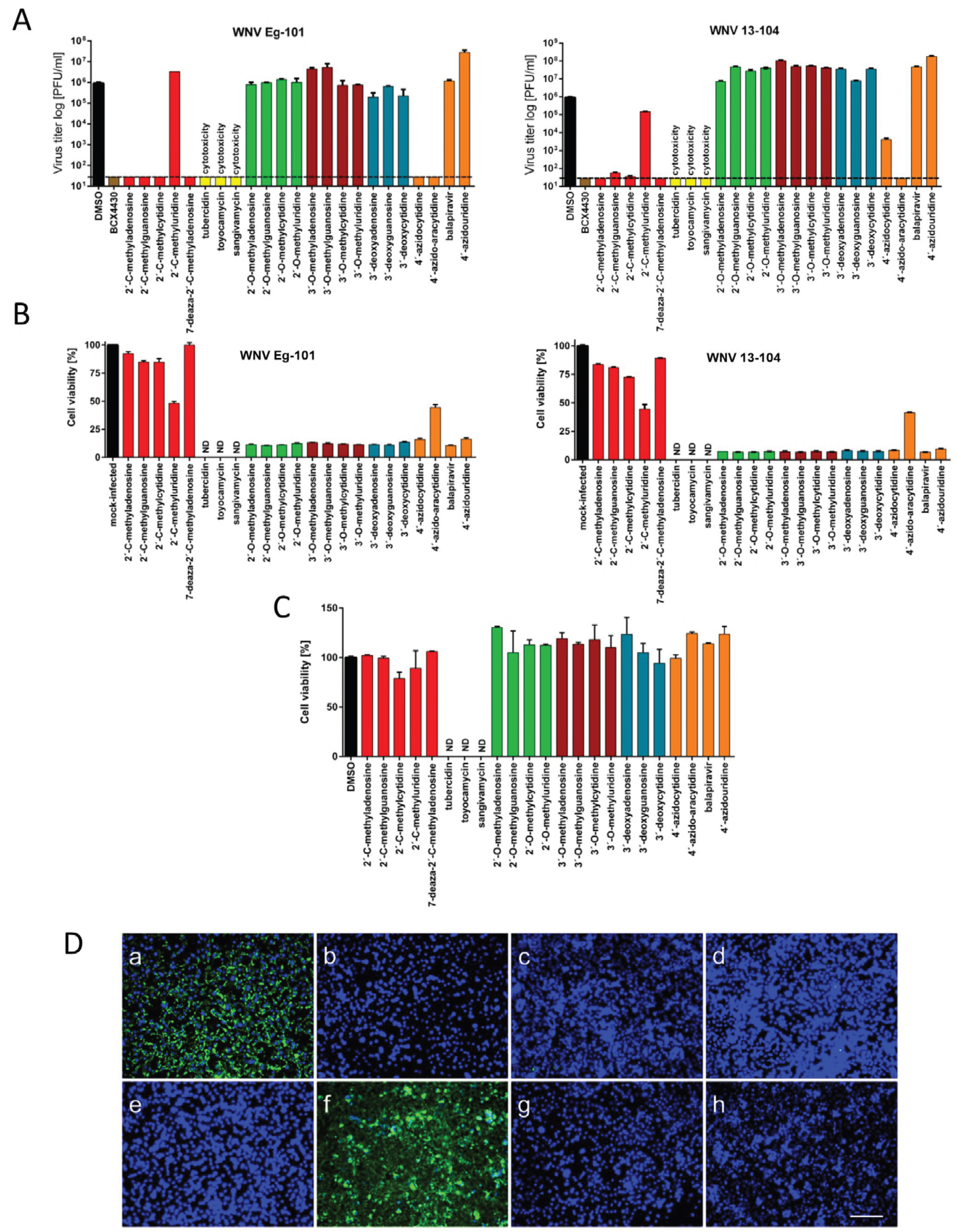
(A) Reduction of West Nile virus (WNV) titers by the indicated nucleoside analogs. PS cells were infected with WNV Eg-101 or 13–104 strain at a multiplicity of infection (MOI) of 0.1 and then treated with 50 µM nucleoside analogs. WNV titers were determined by plaque assay 3 days post-infection. Viral titers are expressed as PFU/mL. (B) Inhibition of WNV-induced CPE by the indicated nucleoside analogs in Vero cells expressed as a percentage of cell viability 3 days post-infection. (C) The cytotoxicity of nucleoside inhibitors was determined by treatment of Vero cells with 50 µM of the indicated nucleoside analogs and was expressed in terms of cell viability 3 days post-infection. Bars indicate the mean values from two independent experiments performed in three replicate wells, and the error bars indicate the standard errors of the means. The horizontal dashed line indicates the minimum detectable threshold of 1.44 log_10_ PFU/mL. ND, not detected (below the detection limit). (D) Inhibition of WNV viral antigen expression by nucleoside inhibitors. PS cells were infected with WNV and treated with 0.5% DMSO (a) or 50 µM 2’-*C*- methyladenosine (b), 7-deaza-2′-CMA (c), 2′-*C*- methylguanosine (d), 2′-*C*- methylcytidine (e), 2′-*C*- methyluridine (f), 4′-azidocytidine (g), or 4′-azido-aracytidine (h). PS cells were fixed with cold methanol:acetone 3 days post-infection and stained with flavivirus-specific antibody labeled with FITC (green) and counterstained with DAPI (blue). Scale bars, 50 µm.

**Figure 2.**
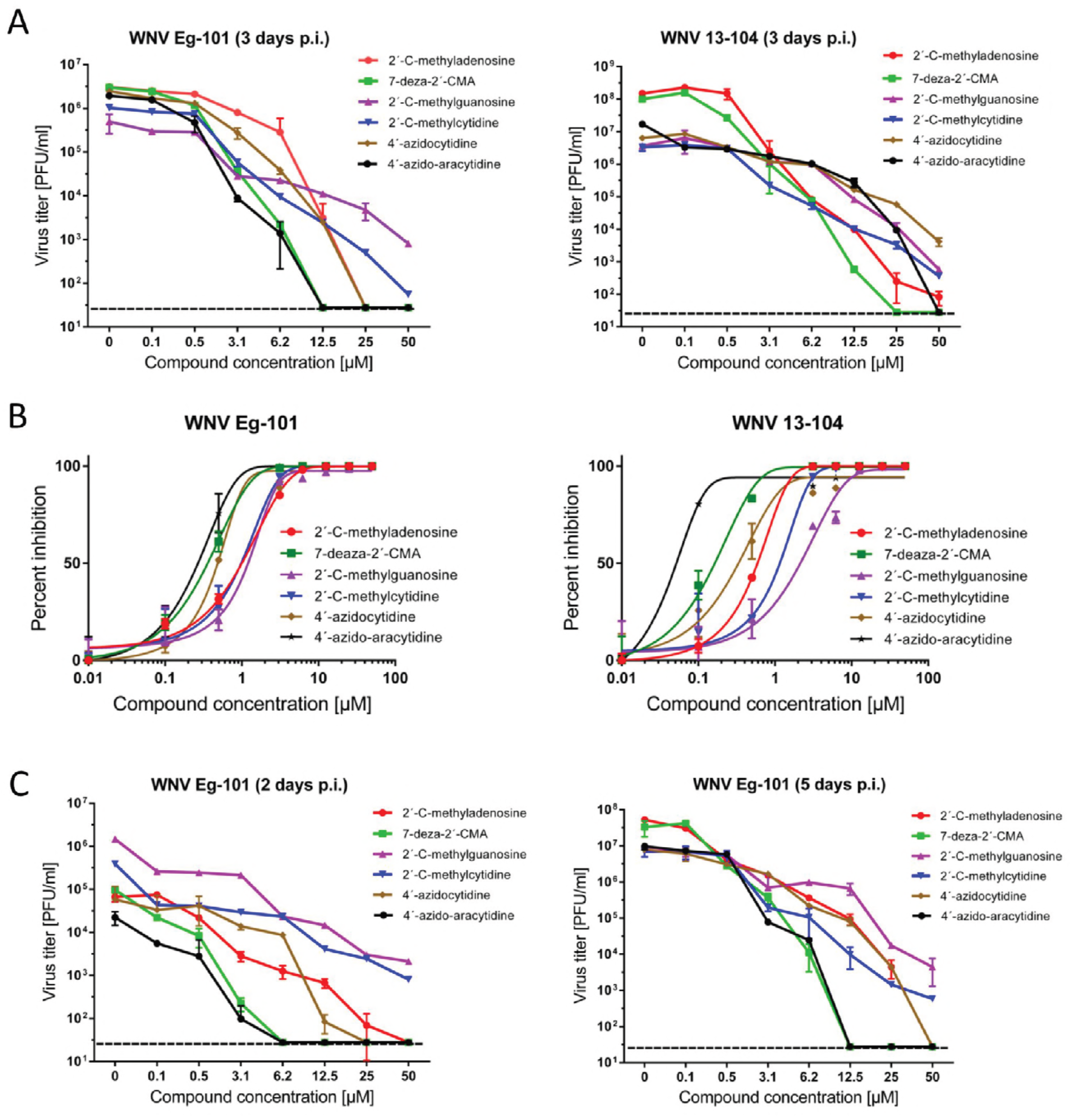
(A) Dose-dependent effects of the indicated West Nile virus (WNV) inhibitors on Eg-101 or 13–104 titer. PS cells were infected with WNV at a multiplicity of infection (MOI) of 0.1 and treated with the appropriate inhibitor at the indicated concentrations 3 days post-infection. The mean titers from two independent experiments done in three biological replicates are shown, and error bars indicate standard errors of the mean. The horizontal dashed line indicates the minimum detectable threshold of 1.44 log_10_ PFU/mL. (B) Sigmoidal inhibition curves for WNV strains Eg-101 and 13–104 in the presence of a serial dilution of indicated nucleoside analogs. (C) Dose-dependent effects of the indicated WNV inhibitors on Eg-101 titer at 2 and 5 days post-infection.

**Table 1.**
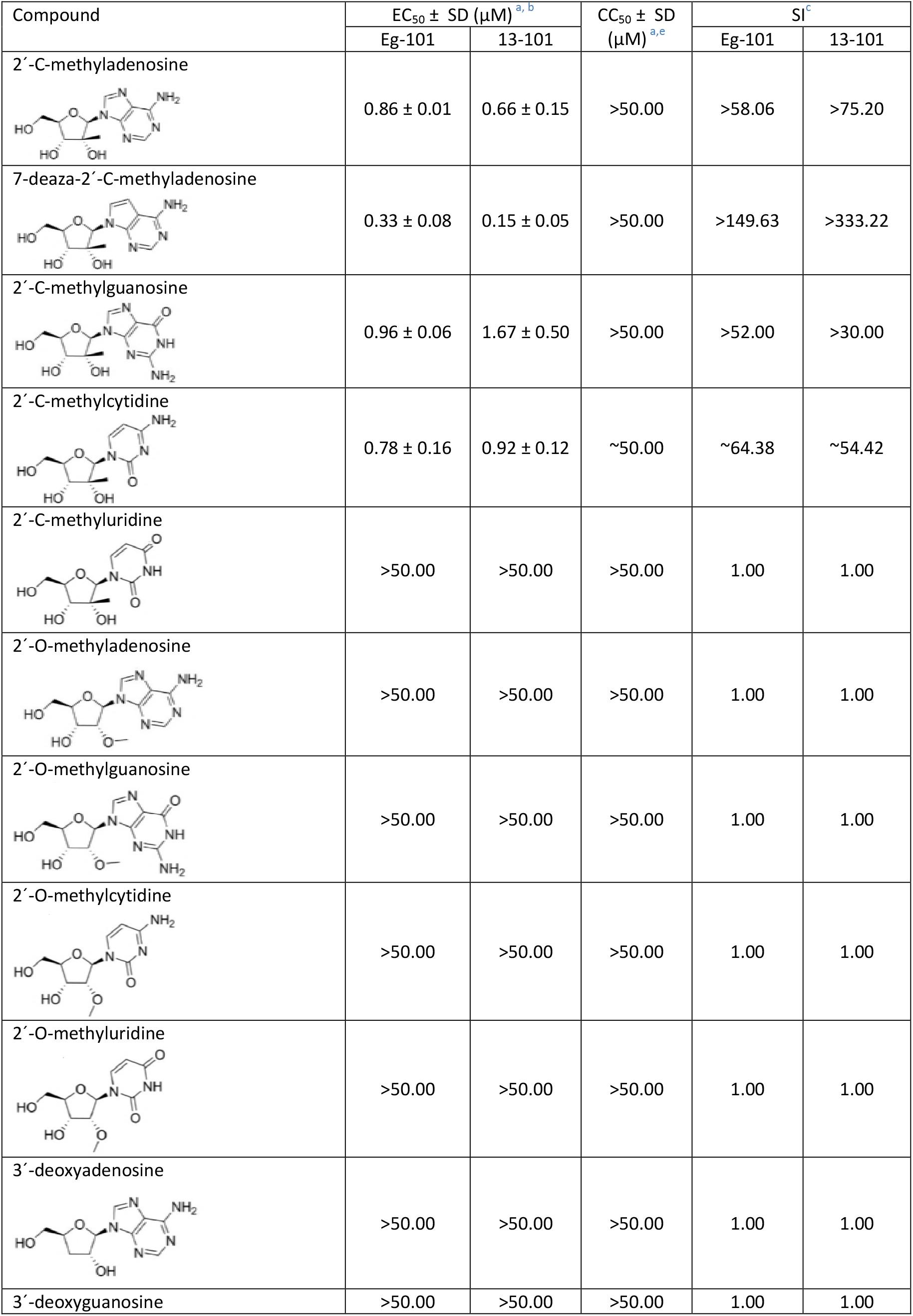
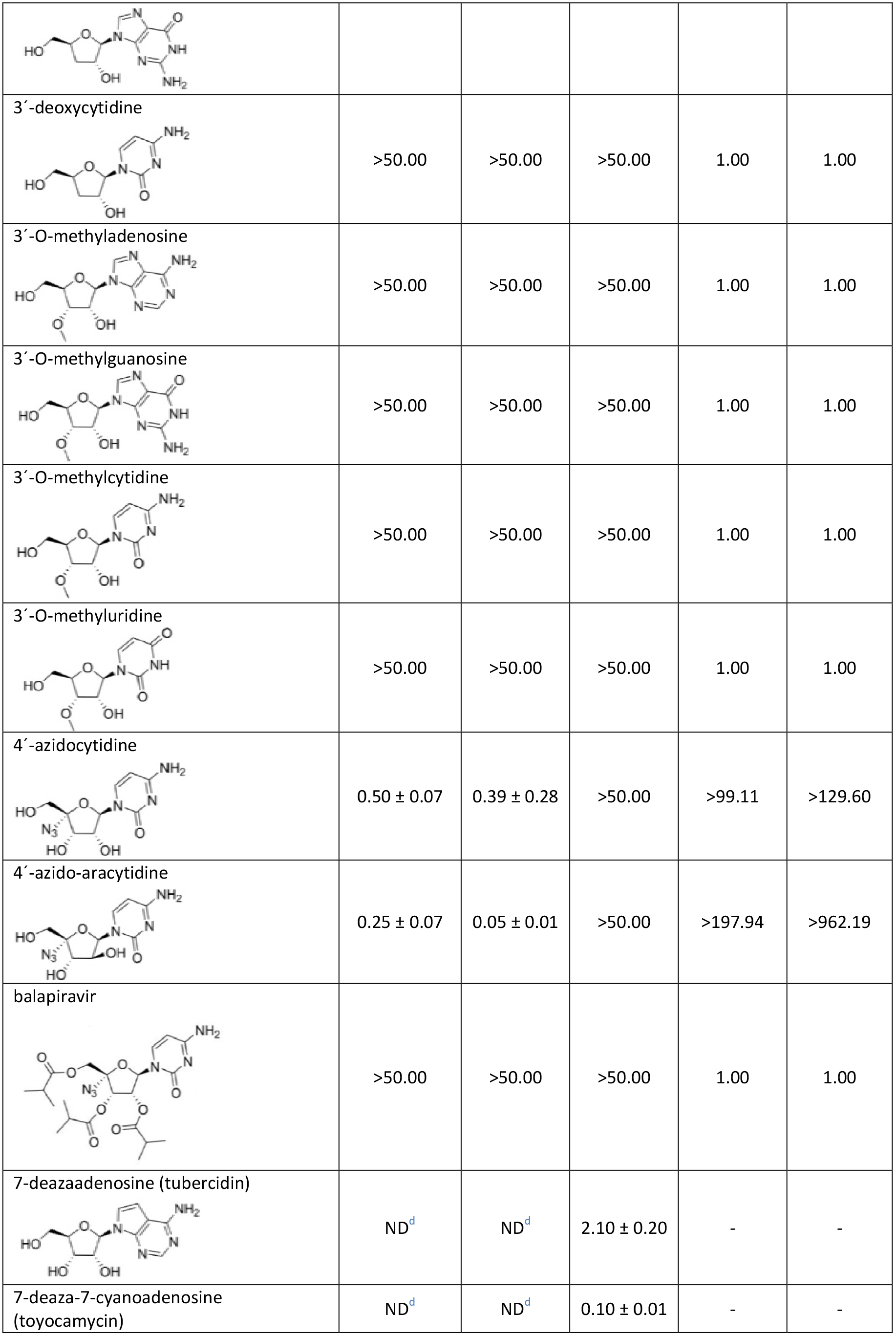
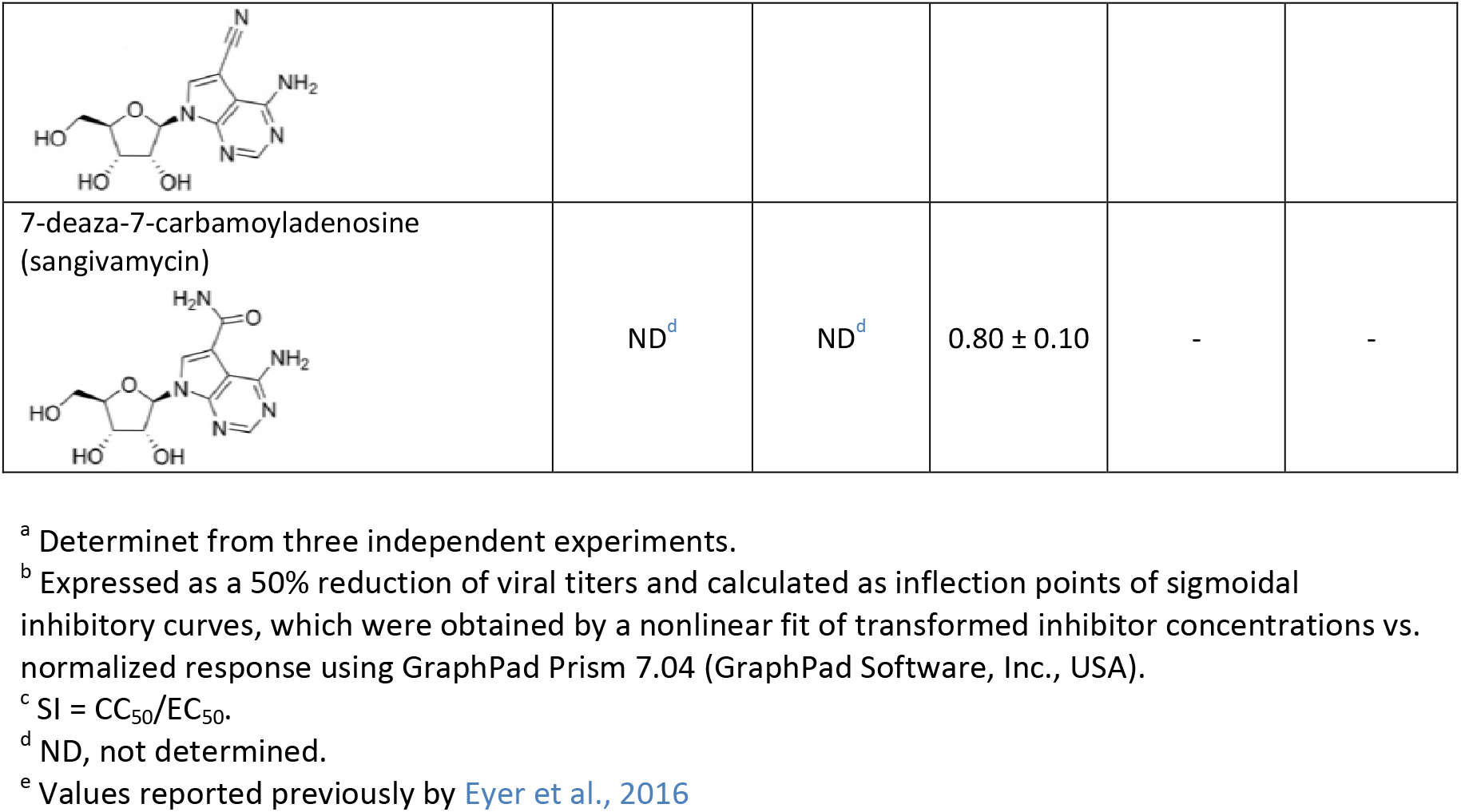
WNV-inhibition and cytotoxicity characteristics of the studied nucleoside analogues

Determinet from three independent experiments.
Expressed as a 50% reduction of viral titers and calculated as inflection points of sigmoidal inhibitory curves, which were obtained by a nonlinear fit of transformed inhibitor concentrations vs. normalized response using GraphPad Prism 7.04 (GraphPad Software, Inc., USA).
SI = CC_50_/EC_50_.
ND, not determined.
Values reported previously by Eyer et al., 2016

2′*-O-*Methyl- or 3′*-O-* methyl-substituted nucleosides demonstrated no inhibitory effects on WNV replication. Similarly, 3′-deoxy nucleosides were completely inactive against WNV (Fig. 1A,B). These compounds did not cause any morphological changes and did not result in decreased PS/Vero cell viability (CC_50_ > 50 µM) at concentrations up to 50 µM after 3 days post-treatment (Table 1). Interestingly, 7-deazaadenosine (tubercidin) and its 7-cyano- and 7-carbamoyl-substituted derivatives (toyocamycin and sangivamycin, respectively) were highly cytotoxic at 50 µM; therefore, their anti-WNV profiles were excluded from further testing (Table 1).

We evaluated four representatives of 4′-azido-modified nucleoside scaffolds for their potential anti-WNV and cytotoxic effects: 4′-azidocytidine (denoted as R-1479), balapiravir (an ester prodrug of 4′-azidocytidine), 4′-azido-aracytidine (a stereoisomeric counterpart of 4′-azidocytidine, denoted as RO-8197), and 4′-azidouridine. 4′-Azidocytidine completely inhibited in vitro replication of the Eg-101 strain at a concentration of 50 µM (EC_50_ value 0.50 ± 0.07 µM) in PS cells. In the case of the 13–104 strain, the inhibitory effect of 4′-azidocytidine was only partial, manifesting as a 10^3^-fold decrease in viral titer at 50 µM compared to mock-infected PS cells. Remarkably strong anti-WNV activity was observed for 4′-azido-aracytidine, with an EC_50_ value of 0.25 ± 0.07 µM for Eg-101 and 0.05 ± 0.01 µM for 13–104 (Fig. 1A). The high antiviral potency of both 4′-azidocytidine and 4′-azido-aracytidine has been demonstrated only in PS cells; in Vero cells the effect was significantly less pronounced (Fig. 1B). The anti-WNV effects of 4′-azidouridine and balapiravir were completely abrogated (EC_50_ > 50 µM; Fig. 1A,B). Though treating the PS cell culture with 50 µM 4′-azido-aracytidine slightly reduced the cell viability (86.9%) (27), the cytotoxicity of the other 4′-azido-substituted nucleosides tested in this study was absent or negligible (Table 1, Fig. 1C).

The antiviral effects of WNV inhibitors identified by viral titer/CPE inhibition assays were confirmed by immunofluorescent staining, which was used to assess the expression of WNV surface E antigen in PS cells as an index of viral infectivity and replication in vitro. Though the surface E protein was highly expressed in WNV-infected mock-treated cells (Fig. 1D), no viral antigen was detected in mock-infected cells (data not shown). Immunofluorescent staining revealed that 50 μM 7-deaza-2′-CMA, 2′-*C*- methyladenosine, 2′-*C-* methylguanosine, 2′-*C*- methylcytidine, or 4′-azido-aracytidine suppressed the expression of WNV surface E antigen, from both the Eg-101 and 13–104 strain, in virus-infected PS cells (Fig. 1D). As expected, WNV surface antigen was expressed extensively in cells treated with 2′-*C*- methyluridine (Fig. 1D). Slight E protein expression was also observed in cells infected with 13–104 strain treated with 4′-azidocytidine (data not shown). The results correlate with the observed inhibition of WNV replication in compound-treated cell cultures.

### Antiviral efficacy of 7-deaza-2′-CMA in a mouse model of lethal WNV infection

Based on 7-deaza-2′-*C*- methyladenosine strongly inhibiting WNV replication in vitro, we used this nucleoside to demonstrate its anti-WNV effect in a mouse model of WNV infection (Fig. 3A). WNV strain Eg101 exhibits low pathogenicity in adult mice (28); therefore, WNV strain 13–104 was used in the in vivo experiments. BALB/c mice infected subcutaneously with a lethal dose of strain 13–104 (10^3^ PFU/mouse) exhibited characteristic clinical signs of infection, such as ruffled fur, hunched posture, tremor, and paralysis of the limbs within days 7 to 12 p.i., with the majority of mice requiring euthanasia (Fig. 3C). The infection was accompanied by a rapid loss of body weight starting 7 days p.i., more than 20% by 12 days p.i. (Fig. 3D). The mortality rate was 95 – 100% with a mean survival time of 10.5 ± 1.9 days (Fig. 2B). Viable virus was detected in the brains of WNV-infected mice 10 days p.i., which was characterized by a mean viral titer of 3.87×10^4^ PFU/mL (Fig. 3E).

**Figure 3.**
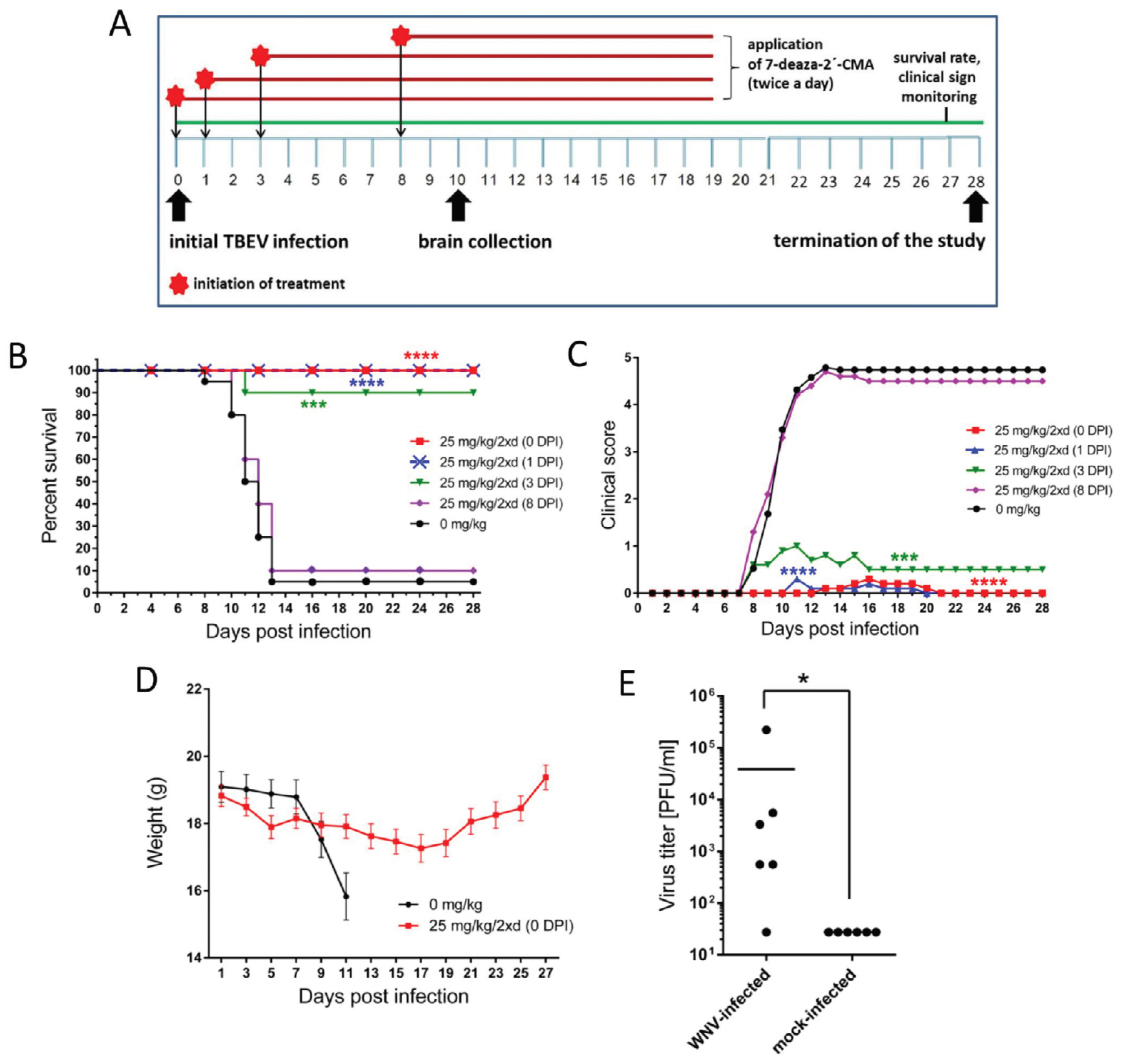
7-Deaza-2′-CMA is effective at treating lethal West Nile virus (WNV) infection in a mouse model. (A) The design of the in vivo antiviral experiment. (B) Groups of adult BALB/c mice were infected with a lethal dose of WNV (strain 13–104) and treated twice daily with intraperitoneal 25 mg/kg 7-deaza-2′-CMA or PBS (mock treatment) at the indicated times after WNV infection. Survival rates were monitored daily for 28 days. (C) Disease signs were scored as follows: 0, no signs; 1, ruffled fur; 2, slowing of activity or hunched posture; 3, asthenia or mild paralysis; 4, lethargy, tremor, or complete paralysis of the limbs; 5, death. (D) Changes in body weight within a 28-day experimental period. Points indicate the mean body weight values of ten mice, and the error bars indicate the standard errors of the means. (E) Groups of adult BALB/c mice were infected with a lethal dose of WNV and treated with 7-deaza-2′-CMA or PBS. On day 10 post-infection, which corresponds with advanced brain infection, the brains of mice were collected and homogenized and the viral titers determined by plaque assay. *P < 0.05; **P < 0.01; ***P < 0.001, ****P < 0.0001.

Treatment with 25 mg/kg of 7-deaza-2′-*C*- methyladenosine twice a day, initiated at the time of virus inoculation and ceased 19 days p.i., resulted in a 100% survival rate (P<0.0001) among WNV-infected mice (Fig. 3B). Nine of ten animals exhibited no clinical signs during the entire 28-day monitoring period; in one mouse we observed slightly bristled fur and slowing activity between 13 and 20 days p.i (Fig. 3C). WNV-infected 7-deaza-2′-CMA-treated mice had a slight loss of body weight (8.2%) by 17 days p.i.; the weight began to increase again after 18 days p.i. (Fig. 3D). No viable virus was demonstrated in the brains of WNV-infected compound-treated mice 10 days p.i. (Fig. 3E). No apparent toxicity or other side effects were observed in mice treated with 7-deaza-2′-CMA (25 mg/kg, twice a day) throughout the treatment period.

A second study was performed to determine the therapeutic effect of treatment initiated at various times after WNV infection. Delayed start of treatment (25 mg/kg, twice daily) 1 day p.i. resulted in a 100% survival rate (P<0.0001; Fig. 3B). Nine of ten animals exhibited no significant signs of disease; in one animal, slightly ruffled fur was observed within 11 to 20 days p.i. (Fig. 3C). Similarly, treatment started 3 days p.i., at the time of peak of viremia, significantly protected the infected animals from disease development and mortality (90% survival rate, P<0.001) and substantially eliminated the appearance of signs of clinical infection (Fig. 3B,C). However, treatment initiated 8 days p.i. (i.e., when the first signs of disease appeared) was inefficient; in this group of animals, characteristic clinical signs of disease were observed within 7 to 12 days p.i. The infection was 90% lethal with a mean survival time of 11.5 ± 1.29 days (Fig. 3B,C).

## Discussion

WNV is a medically important emerging arbovirus able to cause serious neuroinfections in humans, against which no approved antiviral therapy is currently available. Therefore, there is an urgent medical need to develop innovative and effective small molecule-based antivirals to treat patients with WNV infection (14). Here, we evaluated the antiviral activity and cytotoxicity of 2′-*C*- methyl-, 2′*-O-* methyl-, 3′*-O-* methyl-, 3′-deoxy-, and 4′-azido-modified nucleosides. These compounds were originally developed for the treatment of chronic hepatitis C (29–35), and some were later repurposed to suppress viral infections caused by tick- and mosquito-borne flaviviruses (27,32,33,36–41).

Our in vitro antiviral screens revealed that a methyl substituent at the *C*2′nucleoside position is an important structural determinant to suppress the replication of WNV in cell culture. Nucleosides methylated in the *C*2′ position have also been reported to strongly inhibit TBEV (27,36,38), DENV (42,43), ZIKV (37,41,44,45), Alkhurma hemorrhagic fever virus, Kyasanur Forest disease virus, Omsk hemorrhagic fever virus, and Powassan virus (39), as well as numerous representatives of the *Picornaviridae* and *Caliciviridae* families (46–48). This indicates that 2′-*C*- methylated nucleosides are broad-range inhibitors of a large variety of single-stranded, plus-sense RNA viruses. We observed that the anti-WNV activity of 2′-*C*- methylated nucleosides is strongly influenced by the identity of heterocyclic nucleobase and was demonstrated to decrease as follows: 2′-*C*- methyladenosine > 2′-*C*- methylcytidine > 2′-*C-* methylguanosine > 2′-*C*- methyluridine. The surprisingly low in vitro antiviral activity of 2′-*C-* methyluridine was also previously reported for TBEV and ZIKV (EC_50_ of 11.1 ± 0.4 µM and 45.45 ± 0.64 µM, respectively) (27,37). This phenomenon could be ascribed to rapid nucleoside degradation with cellular uridine phosphorylase, a cytoplasmic enzyme playing an important role in uridine catabolism (49,50).

In contrast, 7-deaza-2′-CMA, in which a *C*2′methylated riboside is bound to a 7-deaza-modified adenine, exhibited significantly increased anti-WNV efficacy and low cytotoxicity in both cell lines. This nucleoside has been reported to have greater metabolic stability, better bioavailability, and longer half-life in beagle dogs and rhesus monkeys than its parent compound, 2′-*C*- methyladenosine (33). Moreover, 7-deaza modification introduced into anti-DENV nucleoside 2′-*C*- ethynyladenosine resulted in a dramatic decrease in compound cytotoxicity, as demonstrated in multiple cell-based systems and in vivo (51). Interestingly, elimination of the *C*2′ methyl from 7-deaza-2′-CMA led to extremely high in vitro cytotoxicity of 7-deazaadenosine (tubercidin) and its derivatives sangivamycin and toyocamycin. Our anti-TBEV screens revealed that further modification of 7-deaza-2′-CMA at position 6 (e.g., 6-hydroxyamino or 6-hydrazinyl), position 7 (e.g., 7-nitro, 7-phenyl, 7-fluoro, 7-cyano, 7-ethynyl, or 7-trifluoromethyl), or positions 6 and 7 (e.g., 6-dymethylamino-7-iodo, 6-cyclopropylamino-7-iodo, or 6-morpholino-7-iodo) led to a complete loss of antiviral activity (unpublished results). We can conclude that 7-deaza modification of adenosine combined with a 2′-*C*- methyl (or 2′-*C*- ethynyl) substituent results in higher antiviral activity, decreased toxicity, and improved pharmacokinetic profile; however, 7-deaza modification itself is responsible for a conspicuous increase in cytotoxicity. On the other hand, the lack of antiviral activity of O2′ or O3′-methylated or 3′-deoxy-modified nucleosides could be explained by the absence of 2’- and 3′-hydroxyls positioned in the α-face of the nucleoside scaffold, which are considered to be crucial for specific hydrogen-bonding interactions of the nucleoside molecule with the flaviviral polymerase active site during RNA replication (27,29). Moreover, 2′*-O-* methylcytidine has been observed to be inefficiently phosphorylated by intracellular kinases and/or extensively deaminated/demethylated in various cell cultures (27).

The 4′-azido moiety appears to be another important determinant of highly efficient inhibition of WNV replication in vitro, as exemplified by two cytidine derivatives: 4′-azidocytidine and 4′-azido-aracytidine. The high anti-WNV activity of both compounds has been observed to be strongly cell type-dependent and could be ascribed to differences in compound uptake or the metabolic processing of both nucleosides by individual cell lines (52). Cell type-dependent antiviral activity of 4′-azido-modified nucleosides was also previously reported for TBEV; the compounds were active in PS cells but their activity completely abrogated in human neuroblasts (27). The cell type-dependent antiviral effect can sometimes affect the results of antiviral screens; therefore, multiple clinically relevant cell lines are needed for antiviral testing (52). Similar to 2′-*C*- methyluridine, 4′-azidouridine exhibits no anti-WNV activity, probably because of low metabolic stability and rapid degradation by cellular phosphorylases (49,50).

Based on our finding that 7-deaza-2′-CMA is a strong inhibitor of WNV replication in cell culture, we evaluated the antiviral effect of this compound in a lethal mouse model of WNV infection. A dose of 25 mg/kg 7-deaza-2′-CMA administered intraperitoneally twice a day starting the day of WNV infection resulted in a 100% survival rate. The treated animals exhibited no clinical signs of neuroinfection and no viral titer was determined in mouse brains. The same treatment regimen with 7-deaza-2′-CMA (25 mg/kg, twice daily, starting at the time of infection) was used to treat TBEV-infected BALB/c mice, which had slightly lower survival rates (60%) and worse clinical scores than WNV-infected animals (36).

The treatment started at the time of infection, but does not correlate with potential human drug use, as the therapy is usually initiated during the neurological phase of the disease. Therefore, we tested the therapeutic effect of treatment initiated at various times after WNV infection. Mice infected with WNV have peak viremia between 48 and 72 hours p.i. The virus then crosses the blood-brain barrier and establishes brain infection (53,54). Therefore, treatment was started in mice before viremia (1 day p.i.), during peak viremia (3 days p.i.), and at the time of established brain infection (8 days p.i.). Delayed start of treatment (25 mg/kg, twice daily) 1 day p.i. resulted in the survival of all infected mice. Even the treatment initiated 3 days p.i. (at the time of a peak of viremia) showed a had high antiviral efficacy, protecting 90% of mice from disease development and mortality. However, animals treated after 8 days p.i. (i.e., at the time of extensive brain infection) had only a slightly higher survival rate and negligibly prolonged mean survival time compared to mock-treated mice. The results suggest that treatment with 7-deaza-2′-CMA is highly effective, even when started 1 or 3 days p.i. (i.e., before the virus infects the brain).

Strong in vivo antiviral activity of 7-deaza-2′-CMA has also been demonstrated in other medically important flaviviruses. This compound substantially improves disease outcome, increases survival, and reduces signs of neuroinfection and viral titers in the brains of BALB/c mice infected with TBEV (36). 7-Deaza-2′-CMA also reduces viremia in AG129 mice infected with the African strain of ZIKV (45) and is a potent inhibitor of DENV in a mouse model of dengue viremia (55). Although 7-deaza-2′-CMA has failed in human clinical tests for chronic hepatitis C treatment, probably due to mitochondrial toxicity (56), this compound could still be suitable and safe for short-term therapy of acute flaviviral diseases, including WNV infections (57), and represents one of the most promising candidates for treatment of flaviviral infections to date.

In conclusion, our results demonstrate that nucleosides with a *C*2′ methyl substituent are potent inhibitors of WNV replication in vitro. The leading structure of this group, 7-deaza-2′-CMA, exerted significant anti-WNV activity in a mouse model of WNV infection. This compound protected WNV-infected mice from disease signs/death, even if the treatment was initiated 3 days p.i. Some other substituents were introduced into the *C*2′nucleoside position previously, particularly 2′-*C*- ethynyl in anti-DENV nucleoside NITD008 (51,57,58) or 2′-fluoro-2′-*C*- methyl in the case of sofosbuvir, which inhibits ZIKV in vitro and in rodent models of ZIKV infection (59). A variety of promising modifications at *C*2′ indicate that this nucleoside position provides a large chemical space for possible changes/substitutions to develop efficient nucleoside inhibitors of flaviviral replication. Another important structural modification for efficient WNV inhibition is 4′-azido substitution of cytidine-based nucleosides. Although these compounds have been tested only in vitro, 4-azido-aracytidine had a nanomolar anti-WNV effect and good cytotoxicity profile. This molecule will be tested in vivo in our future studies. Our data strongly suggest that nucleoside analogs represent a rich source of promising inhibitors for the further design and development of innovative and effective drugs active against important human pathogens, including flaviviruses.

## Materials and methods

### Ethics statement

This study was carried out in strict accordance with Czech law and guidelines for the use of experimental animals and protection of animals against cruelty (Animal Welfare Act 246/1992 Coll.). All procedures were reviewed by the local ethics committee and approved by the Ministry of Agriculture of the Czech Republic (permit no. 22006/2016-MZE-17214).

### Cell cultures, virus strains, and antiviral compounds

PS cells (60), a cell line widely used for the isolation and multiplication of arthropod-borne flaviviruses, were cultured at 37°C in L-15 (Leibovitz) medium containing 1 mM L-glutamine, 4% fetal bovine serum, and 1% penicillin and streptomycin (Sigma-Aldrich, Prague, Czech Republic). Epithelial kidney (Vero) cells from *Cercopithecus aethiops* were cultured in MEM (Minimum Essential Medium) containing 10% fetal bovine serum, 5% L-glutamine, and 1% penicillin and streptomycin (Sigma-Aldrich, Prague, Czech Republic) at 37°C in a 5% CO_2_ atmosphere. Two distinct low-passage WNV strains were used to evaluate the anti-WNV activity of the tested compounds. Eg-101, a member of genomic lineage 1, was originally isolated from human serum in Egypt in 1951 (61). WNV strain 13–104, a representative of genomic lineage 2, was isolated from the *Culex modestus* mosquito in the Czech Republic in 2013 (62).

2′-*C*-Methyl-, 2′*-O-* methyl-, and 3′*-O-* methyl-substituted nucleoside analogs and 3′-deoxynucleosides were purchased from Carbosynth (Compton, UK). 4′-Azido-modified nucleosides and BCX-4430 were obtained from Medchemexpress (Stockholm, Sweden). Tubercidin, toyocamycin, and sangivamycin were purchased from Sigma-Aldrich (Prague, Czech Republic). 7-Deaza-2′-*C*- methyladenosine was synthesized at the Institute of Organic Chemistry and Biochemistry in Prague. Nucleoside analogs were diluted in 100% dimethylsulfoxide (DMSO) to 10 mM stock solutions.

### Antiviral assays

In order to perform the viral titer reduction assay, PS cells were seeded in 96-well plates (approximately 3 × 10^4^ cells per well) and incubated for 24 hours at 37°C to form a confluent monolayer. The medium was then aspirated from the wells and replaced with 200 µl of fresh medium containing 50 µM of the test compounds and appropriate virus strain at a multiplicity of infection (MOI) of 0.1. Virus-infected cells treated with BCX-4430 (63) and DMSO (mock-treated cells) were used as positive and negative controls, respectively. The culture medium was collected 3 days p.i. and viral titers (expressed as PFU/mL) determined from the collected culture medium by plaque assays (38,64).

As the antiviral effect of many compounds is cell-type dependent, we further tested the anti-WNV activity of nucleoside analogs in Vero cells in cytopathic effect (CPE) inhibition assays. Vero cells were seeded in 96-well plates, incubated to form a confluent monolayer, and treated with virus and tested compounds as described for the viral titer inhibition assay. The cells were examined microscopically for virus-induced CPE. Three days p.i., the supernatant medium was aspirated from the cells and the cell culture stained with naphtalene black. The rate of CPE was expressed in terms of cell viability as the absorbance at 540 nm by compound-treated cells relative to the absorbance by DMSO-treated cells.

The dose-dependent antiviral activities of selected WNV inhibitors were assessed as follows. PS cells were seeded in 96-well plates (approximately 3 × 10^4^ cells per well) and incubated at 37°C to form a confluent monolayer. After a 24-hour incubation, the medium was aspirated from the wells and replaced by a fresh medium containing the tested compounds over the concentration range of 0 to 50 µM and appropriate WNV strain (MOI 0.1). The culture medium was collected continuously 2, 3, and 5 days p.i. The viral titers (expressed as PFU/mL) were determined from the collected supernatant media by a plaque titration assay and used to construct dose-dependent and inhibition curves. The viral titers obtained 3 days p.i. were used for calculation of the 50% effective concentrations (EC_50_; the concentration of compounds required to inhibit the viral titer by 50% compared to the control value).

### Immunofluorescent staining

The results obtained from antiviral assays were confirmed by a cell-based flavivirus immunostaining assay, a method based on determining the compound-induced inhibition of viral surface antigen (E protein) expression (38). Briefly, PS cells seeded on 96-well plates were treated with the test compound (50 μM) and infected with WNV strains Eg-101 or 13–104 at an MOI of 0.1. After a 3-day incubation at 37°C, the cell monolayers were fixed with cold acetone-methanol (1:1), blocked with 10% fetal bovine serum and incubated with a mouse monoclonal antibody that specifically recognizes the flavivirus group surface antigen (1:250, Sigma-Aldrich, Prague, Czech Republic). After extensive washing, the cells were labeled with an anti-mouse goat secondary antibody conjugated with FITC (1:500) and counterstained with DAPI (1 μg/mL) to visualize the cell nuclei. The fluorescence signal was recorded with an Olympus IX71 epifluorescence microscope.

### Cytotoxicity assay

PS or Vero cells grown in 96-well plates for 24 hours to form a confluent monolayer were treated with test compounds over the concentration range of 0 – 50 µM. Three days post-treatment, the cell culture medium was collected and the potential cytotoxicity of test nucleosides determined in terms of cell viability using Cell Counting Kit-8 (Dojindo Molecular Technologies, Munich, Germany) according to the manufacturer’s instructions. The compound concentration that reduced cell viability by 50% was considered the 50% cytotoxic concentration (CC_50_).

### Mouse infections

To evaluate the anti-WNV effect of 7-deaza-2′-CMA in vivo, five groups of 6-week-old female BALB/c mice (purchased from AnLab, Prague, Czech Republic) were subcutaneously injected with WNV strain 13–104 (1000 PFU/mouse) and treated intraperitoneally with 200 µL of 7-deaza-2′-CMA (25 mg/kg twice a day at 8-hour intervals) as follows: group 1 (n = 10), treatment initiated immediately after infection (0 days p.i.); group 2 (n = 10), treatment initiated 1 day p.i.; group 3 (n = 10), treatment initiated 3 days p.i., group 4 (n = 10), treatment initiated 8 days p.i. (when the first clinical signs appeared); and group 5 (n = 10), control animals, treated with vehicle only. 7-Deaza-2′-CMA was freshly solubilized in sterile saline buffer before each injection and administered to the animals for 19 (group 1), 18 (group 2), 17 (group 3), and 11 (group 4) days. The clinical scores, body weight, and survival rates of WNV-infected mice were monitored daily over 28 days. Illness signs were evaluated as follows: 0 for no symptoms; 1 for ruffled fur; 2 for slowing of activity or hunched posture; 3 for asthenia or mild paralysis; 4 for lethargic, tremor, or complete paralysis of the limbs; 5 for death. All mice exhibiting disease consistent with clinical score 4 were terminated humanely (cervical dislocation) immediately upon detection.

For determination of the WNV titer in mouse brains, a group of 6-week-old BALB/c mice were infected subcutaneously with 1000 PFU of TBEV and immediately treated with 7-deaza-2′-CMA twice daily at a concentration of 25 mg/kg. Ten days p.i., when the clinical symptoms of WNV disease were clearly observable in mice treated with vehicle only, the animals were sacrificed and the brains collected, weighed, homogenized using Precellys 24 (Bertin Technologies), and prepared as 20% (w/v) suspensions in saline. Each homogenate was centrifuged at 5000 × *g* and the supernatant medium used to determine virus titer by plaque assay.

### Statistical analysis

Data are expressed as mean ± standard deviation (SD), and the significance of differences between groups was evaluated by the Mann-Whitney *U* test or ANOVA. Survival rates were analyzed using the log rank Mantel-Cox test. All tests were performed using GraphPad Prism 7.04 (GraphPad Software, Inc., USA). P<0.05 was considered significant.

## Acknowledgments

This study was supported by a grant from the Czech Science Foundation (grant 16–20054S). The funder had no role in the study design, data collection and analysis, decision to publish, or preparation of the manuscript.

